# Examination of the efficacy of small genetic panels in genomic conservation of companion animal populations

**DOI:** 10.1101/2020.02.04.934158

**Authors:** Aaron J. Sams, Brett Ford, Adam Gardner, Adam R. Boyko

## Abstract

In many ways dogs are an ideal model for the study of genetic erosion and population recovery, problems of major concern in the field of conservation genetics. Genetic diversity in many dog breeds has been declining systematically since the beginning of the 1800’s, when modern breeding practices came into fashion. As such, inbreeding in domestic dog breeds is substantial and widespread and has led to an increase in recessive deleterious mutations of high effect as well as general inbreeding depression. Pedigrees can in theory be used to guide breeding decisions, though are often incomplete and do not reflect the full history of inbreeding. Small microsatellite panels are also used in some cases to choose mating pairs to produce litters with low levels of inbreeding. However, the long-term impact of such practices have not been thoroughly evaluated. Here, we use forward simulation on a model of the dog genome to examine the impact of using limited markers panels to guide pairwise mating decisions on genome-wide population level genetic diversity. Our results suggest that in unsupervised mating schemes, where breeding decisions are made at the pairwise-rather than population-level, such panels can lead to accelerated loss of genetic diversity compared to random mating at regions of the genome unlinked to panel markers and demonstrate the importance of genome-wide genetic panels for managing and conserving genetic diversity in dogs and other companion animals.

## INTRODUCTION

Loss of genetic diversity in small populations is a major concern of the global conservation community because it can reduce fitness and adaptability, and can ultimately lead to breed, population, or species extinction (O’Grady et al., 2006). For example, Target 13 of the Convention on Biological Diversity’s Strategic Plan (https://www.cbd.int/sp/) is aimed at “minimizing genetic erosion” of both socio-economically and culturally valuable species and “safeguarding their genetic diversity.’’

It has been repeatedly demonstrated that the optimal way to reduce genetic erosion in management programs is to minimize the mean kinship within a population (Ballou & Lacy, 1995; Fernández, Toro, & Caballero, 2001; Sonesson & Meuwissen, 2001). Kinship can be estimated using pedigrees, however, pedigrees are frequently inaccurate, incomplete, or missing (Cassell, Adamec, & Pearson, 2003). In lieu of deep, high quality pedigrees, molecular data is used. While there has been a push in recent years to move towards monitoring and managing populations of conservation concern with genomic data in the form of whole genome genotyping or whole genome sequencing (Flanagan, Forester, Latch, Aitken, & Hoban, 2017; Ivy, Putnam, Navarro, Gurr, & Ryder, 2016; Leroy et al., 2017; Shafer et al., 2015), many populations are still monitored with small sets of neutral markers, typically tens to hundreds of microsatellites (Abdul-Muneer, 2014; Attard et al., 2016; Kaczmarczyk, 2016; Kirk & Freeland, 2011; Pedersen, Pooch, & Liu, 2016; Song et al., 2018; Toro, Fernández, & Caballero, 2009).

While many researchers have cautioned against using small marker panels alone to guide captive breeding (see for example (Miguel Toro et al., 1998; Wang & Hill, 2000)) at least some previous research has suggested that small microsatellite panels can be used to maintain genetic diversity in captive breeding programs (Kaczmarczyk, 2016). Importantly though, as Nicholas and colleagues (*Nicholas, Mellersh, & Lewis, 2018) recently pointed out, such small microsatellite panels do not effectively survey genome-wide genetic diversity. Rather, they survey genetic diversity at and near (depending on the extent of linkage disequilibrium) the assayed microsatellites. Direct comparison of microsatellite panel and genome-wide (e.g. single nucleotide polymorphisms (SNPs)) panel estimates of genetic diversity are rare, but the argument of Nicholas and colleagues is supported by a recent direct comparison in *Arabidopsis halleri*, which found that microsatellite-based estimates of genetic diversity and population differentiation differ substantially from unbiased estimates from SNPs (Fischer et al., 2017).

In many ways dogs are an ideal model for the study of genetic erosion and population recovery. Genetic diversity in many common domestic dog breeds has been declining systematically since the beginning of the 1800’s, when modern breeding practices came into fashion (Jansson & Laikre, 2018). As such, inbreeding in domestic dog breeds is substantial and widespread (Freedman et al., 2014; Kettunen, Daverdin, Helfjord, & Berg, 2017; Pedersen, Pooch, & Liu, 2016; Sams & Boyko, 2018) and has led to an increase in recessive deleterious mutations of high effect (Jagannathan et al., 2019; Marsden et al., 2016) as well as general inbreeding depression (Chu et al., 2019).

Dog breeders and breed clubs are increasingly aware of the serious consequences of diversity loss, and with robust panels of both microsatellite markers and genome-wide SNP arrays widely available commercially, there is great potential for breeders to use genetic testing in ways that ultimately improve (or worsen) genetic diversity. However, a key challenge, at least in the United States, is a lack of population-level management. Rather, individual dog breeders or groups of breeders typically manage small subsets of a breed, often relying on pedigrees or commercially available molecular tests to minimize known genetic health risks and sometimes overall inbreeding in individual litters of dogs. For breeds still managed in such a way, it is critical to long-term breed health and survival to understand the long-term impacts on genome-wide genetic diversity of chosen mating strategies and the molecular tools used to guide those strategies.

As a first step in understanding the impact of such population management, we conducted individual-based forward-time population genetic simulations of linked genetic diversity on a model of the dog genome using SLiM 3 (Haller & Messer, 2019). We apply a range of human-directed mate choice models to ask how well different mate choice schemes applied to a restricted panel of 33 “microsatellite” locations in our model dog genome (referred to throughout as MS33) affect genetic diversity genome-wide. More specifically, we evaluate several combinations of metrics calculated on this set of 33 multi-allelic markers to guide diversity-based mate choice including heterozygosity, internal relatedness (IR) (Amos et al., 2001) and average genetic relatedness (AGR) (Wang, 2002). Importantly, we do not model the generally recommended strategy of selecting parents to minimize population-level kinship, but rather simulate individual mating decisions aimed at optimizing genetic measurements for single offspring. We ask how mate choice using these methods impacts genome-wide genetic variation as reflected by heterozygosity and allelic richness (average number of alleles per locus) compared to random mate choice, as well as mate choice guided by two generation pedigree awareness, genome-wide heterozygosity, and relatedness calculated from genome-wide identity by descent (see Materials and Methods).

## MATERIALS AND METHODS

### Genome Model

We implemented all simulations in SLiM (v3.3) (Haller & Messer, 2019) and all simulations were run in parallel using Amazon Web Services EC2, SQS, and auto-scaling services. The genome model in our simulations is a rough approximation of the canine genome. For computational efficiency, we model genetic variation as non-recombining 0.5 megabase multi allelic haplotype blocks. These haplotype blocks are represented by mutation type 1 (m1) in the simulation template (Appendix S1). Chromosomes are created by dividing the genome into 38 sets of 120 haplotype blocks, approximating a genome size of 2.28 gigabases. Recombination rate between chromosomes is 0.5 and within chromosomes is 0.005, based on an overall mutation rate per base-pair of 10^−8^. Additionally, we modeled 33 microsatellite loci (m2 in the simulation template) spaced across the first 25 chromosomes as such: 3 on chromosome 1, 2 each on chrs 2 — 7, 1 each on chrs 8 — 25. This distribution of markers across chromosomes is similar to the 33 STR panel used in (Pedersen, Pooch, & Liu, 2016). Given a number of unique haplotype blocks and microsatellite alleles at the start of the simulation burn-in, we evenly distributed those alleles across individuals in the founding population (see Appendix S2 — Figure S2 for a graphical example of this model).

### Demographic Model

We created a relatively simple demographic model in which a single ancestral population evolves for a burn in and drift period to allow founding genetic diversity to recombine sufficiently, and experience sufficient genetic drift. This is followed by a short immediate bottleneck. Finally, the population expands and goes through a number of mate-choice generations. Population genetic data is collected at the beginning of the first generation of mate choice, and again at the very end of the simulations (see Appendix S2 — Figure S3 for a graphical example of this model).

### Life Cycle Model

Generations in the initial burn in/drift period and bottleneck follow the standard Wright-Fisher model in SLiM. However, during mate-choice, we induce a slightly different model. Creation of offspring in the mate-choice schema includes (see Appendix S2 — Figure S4 for a graphical example of this model):

1. Random sampling of a parent from the full population (sample individual that has not been mated > MAXIMUM_NUMBER_OF_MATINGS times).
2. Choose second parent:
  a. Sub-sampling a fraction of the remaining population randomly (according to MATING_POOL_SIZE) from which to choose potential mates. This step is intended to model the fact that within an effective population of purebred dogs, individual dogs only have access to a limited number of other dogs as mates. This sampling is repeated if no eligible (has been mated < MAXIMUM_NUMBER_OF_MATINGS times) dogs are sampled.
  b. A mate for the first individual is chosen according to a specific mate choice model (see Mate Choice Models) from this pool.
3. In the event of a layered mate-choice model in which two statistics are used to choose mates, a PROPORTION_OF_MATES_FOR_LAYERED_MATE_CHOICE parameter is used to sub-sample the potential mates based on each statistic used.

### Mate Choice Models

Mate choice models that we implemented in this study primarily differ in how a second parent is selected. First parents are selected randomly from the entire population as described above.

#### Random

Second parent is randomly selected from the sampled mating pool.

#### Pedigree

Second parent is the individual with the lowest relatedness as calculated from three generation pedigrees, randomly sampled in ties. In other words, using the keepPedigrees=True option in SLiM 3, pedigree relatedness between individuals in the current generation can be calculated from pedigrees. This option maintains pedigrees including all current individual’s parents and grandparents.

#### Heterozygosity models [Microsatellite (MS33-HET) and Genome-wide (GW-HET)]

For these models we calculate the expected heterozygosity for offspring between the first parent and all individuals in the mating pool as the average pairwise observed homozygosity across all four pairwise combinations of parental genomes (assuming no recombination). We choose as the best mate the individual that would produce offspring with the highest heterozygosity and select randomly amongst ties.

#### Internal Relatedness (MS33-IR)

For this model, as above, we calculate mean Internal Relatedness (Amos et al., 2001) amongst four possible gametic pairs at each microsatellite locus and then average across all loci and choose the individual that produces the lowest IR value, selecting randomly from ties.

#### Average Genetic Relatedness (MS33-AGR)

Here we apply a method designed to calculate relatedness between individuals based on small panels of SNPs or microsatellites (Wang, 2002). We based the code in our SLiM template on the implementation of this calculation found in the R package Demerelate (https://github.com/cran/Demerelate) and ran several tests to ensure that our SLiM implementation and the R version produced identical results. We chose as the second parent the individual that produced the lowest AGR value, selecting randomly from ties.

#### Whole Genome Relatedness (GW-REL)

We calculate whole genome relatedness as in (Hedrick & Lacy, 2015) and as above, calculated relatedness across all four possible gametic pairs between two individuals. We chose as the second parent the individual least related to the first randomly selected parent and chose amongst ties randomly.

#### Layered Mate Choice (MS33_IR_AGR)

We additionally investigated a single microsatellite mate choice model combining the IR and AGR statistics. First, a fraction of individuals in the mating pool are chosen based on the IR statistic, and then an individual is chosen from that sample based on AGR (see Life Cycle Model above).

### Population Genetic Statistics

#### Observed Heterozygosity

We calculated per individual as the fraction of all genotypes in an individual that are heterozygous. In some cases we averaged observed heterozygosity across all individuals in a population.

#### Allelic Richness

We calculated allelic richness as the total number of unique alleles at each position. In some outputs (results not presented here but data available) we also calculated richness for alleles >= 0.05 frequency.

#### Coefficient of inbreeding

We calculated the coefficient of inbreeding (COI) for individual dogs as the fraction of all mutations of type ‘m1’ that are identical within an individual divided by the total genomic length.

### Statistical Analyses

For nearly all statistical comparisons, including calculations of 95% confidence intervals used in Figure 1 we have utilized estimation graphics and statistics (Ho, Tumkaya, Aryal, Choi, & Claridge-Chang, 2019) as implemented in the python package dabest (v0.2.5 - https://acclab.github.io/DABEST-python-docs/index.html). See python script used to generate results for specific uses (github link will be added upon acceptance).

**Figure 1.**
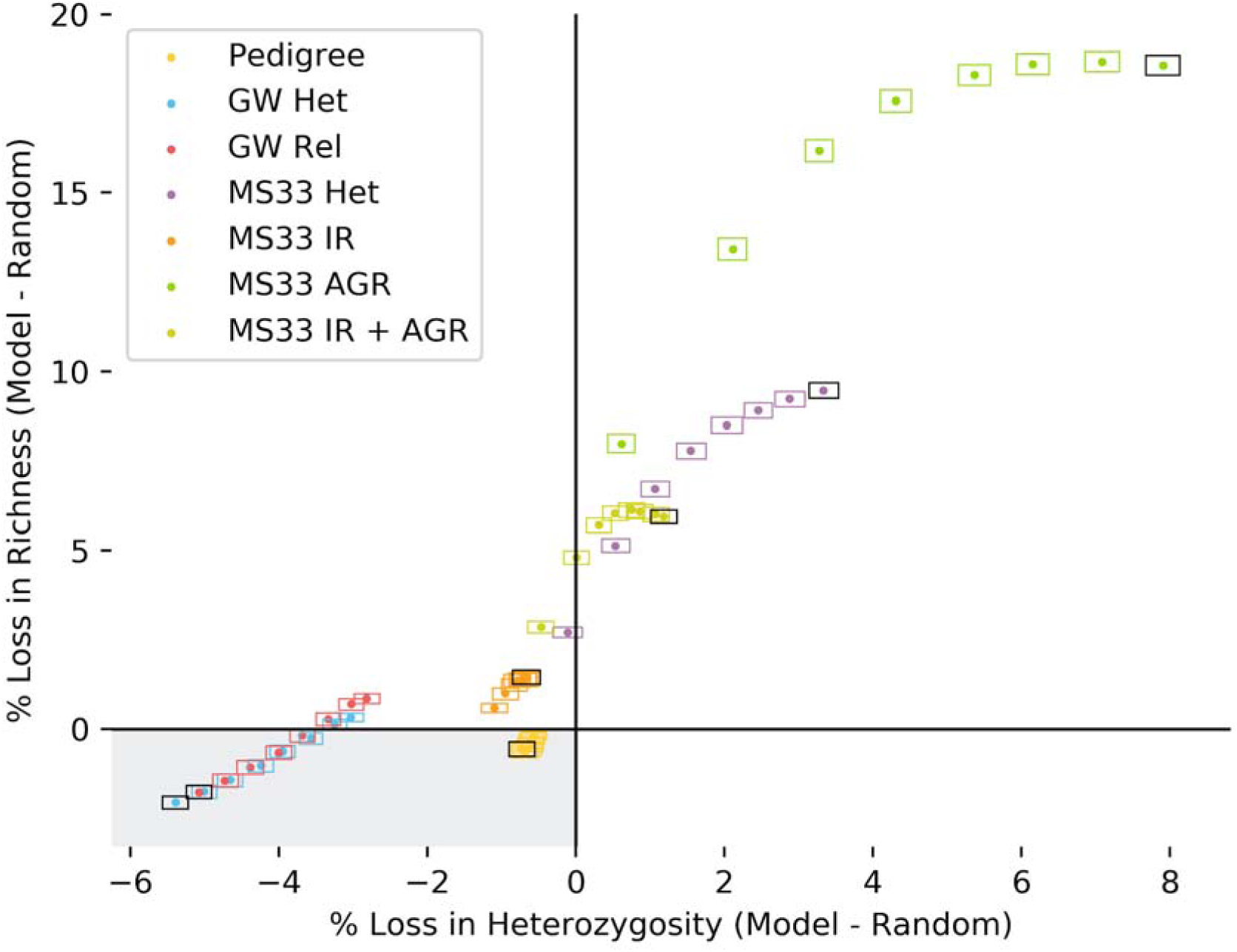
Small marker panel mate selection performs worse and genome-wide marker panel mate selection improves over time relative to random mating. Illustration of the percent loss in heterozygosity and richness over the duration of simulations for parameter set 1, relative to loss during random mating simulations. Dots represent the mean difference between each mate choice model and random mating using a randomization method. Height and width of boxes represent 95% confidence intervals for richness and heterozygosity, respectively (see Materials and Methods). Black boxes represent the mean difference over the duration of simulations (40 generations of mate choice). Grey box represents the quadrant in which both loss of heterozygosity and richness is slowed relative to random mating. GW models (including Pedigree) improve over the course of simulations, while MS models lose more diversity over time than random mating.

## RESULTS

### Genetic variation in simulated populations compared to present-day dogs

For each of eight mate choice models (7 described in Materials and Methods and an additional compound mate choice model of MS33-IR and MS33-AGR), we generated 100 replicate simulations for each of 21 different demographic parameter sets. Three baseline parameter sets vary in the effective population size used during the burn in period. Additional simulations are variants of these three baseline parameters, varying one other parameter (Appendix S3 — Table S1).

The simulated populations show realistic genetic variation compared to present-day domestic dog breeds. For example, across all models, prior to mate selection the mean coefficient of inbreeding (COI) across all 800 replicates (100 replicates x 8 model types) for parameter set 1 (PS-1) is approximately 0.17 — 0.21, and across all simulations is ∼0.12 — 0.38, well within the range of several common dog breeds today (Sams & Boyko, 2018). Similarly, mean internal relatedness (IR) varies in the range of -0.013 — 0.034 across PS-1 and -0.015 — 0.1 across all parameter sets (See Appendix S3 — Tables S2 and S3 for COI and IR means by model and parameter set). Pedersen and colleagues (Pedersen, Pooch, & Liu, 2016) observed a mean IR value of 0.007 in 102 Bulldogs.

### MS33-based mate-choice models lead to more diversity loss than random mate-choice or short pedigrees over time

Performance of all models was measured as the loss of genetic diversity relative to the loss of genetic diversity observed across the random mating model. Of the MS33-based models tested, Internal Relatedness-based mate choice (MS33-IR) performed best overall after 40 generations of mate choice. This model limited loss of heterozygosity comparably or slightly better than random mating across all parameter sets tested (Figure 1, Appendix S3 — Table S4). However, this model performed worse than random mating at preserving allelic richness across all parameter sets but one (PS-19) at which it was indistinguishable from random mating (Appendix S3 — Table S5).

With a few exceptions, all other MS33-based models performed worse than random mating using both heterozygosity and allelic richness as a metric of diversity loss. In these cases, diversity loss is as great or in some cases substantially greater than random mate choice and recent pedigree-based mate choice. Importantly, even in cases where MS models preserve heterozygosity more than random mating, the trajectory of diversity loss over time in these simulations suggests that given enough time these models would also perform worse overall than random mating (Figure 1, Appendix S2 — Figure S1).

Among the MS33-based models, MS33-IR mate choice lost the least amount of genetic diversity, and MS33-AGR mate choice lost the greatest amount of genetic diversity. In all cases, the accelerated loss of genetic diversity compared to random mating is due to preserving diversity at a small number of loci at the expense of the remainder of the genome. In other words, by avoiding inbreeding with individuals more closely related at a small number of loci scattered throughout the genome, the effective population size at unlinked loci, which are evolving under drift, is further reduced.

As direct evidence of this reduction of effective size at unlinked loci, we observe that genetic diversity at MS loci is preserved well, but is lost more than random mating away from these MS positions (Figure 2A). This pattern is consistent regardless of the MS-based mate-selection model we examine and is not observed in genome-wide models described below (Figure 2B).

**Figure 2.**
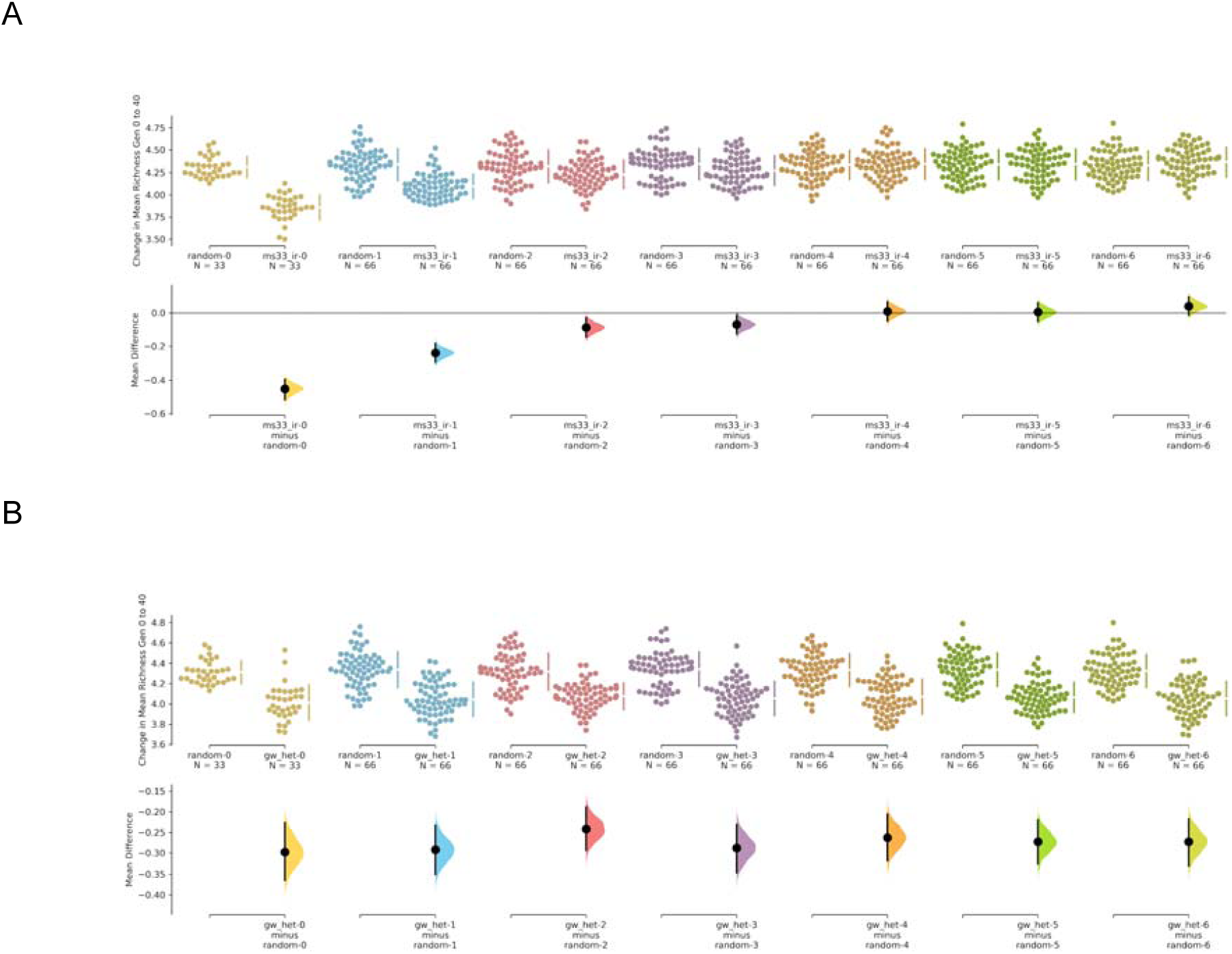
Local preservation of allelic richness degrades with distance to MS markers in MS but not GW mate choice models. Mean difference in percent allelic richness lost during 40 generations of mate choice between Random and other models. A) In MS models such as MS33-IR, richness is better preserved near markers used during mate choice but degrades with distance. B) In GW models like GW-HET, richness is consistently preserved across all markers.

### Number of repeated matings can moderate “popular parent” effects

We examined whether adjusting the total number of individuals that any single individual can access as potential mates within the population, termed here “mating pool size,” as well as the maximum number of times any single individual can contribute to the next generation, termed here “number of repeated matings,” has a substantial impact on the magnitude of the loss of diversity in MS-based mate selection models.

We reduced the maximum number of matings per individual per generation from 200 (the population size) to 50 (PS10-12) and 5 (PS13-15). Reducing from 200 to 50 had no significant impact on diversity metrics (Figure 3A, Appendix S3 — Tables S6-8). However, reducing from 200 to 5 led to a substantial reduction in loss of heterozygosity and allelic richness, and lower increase in COI (Figure 3B, Appendix S3 — Tables S6-8).

**Figure 3.**
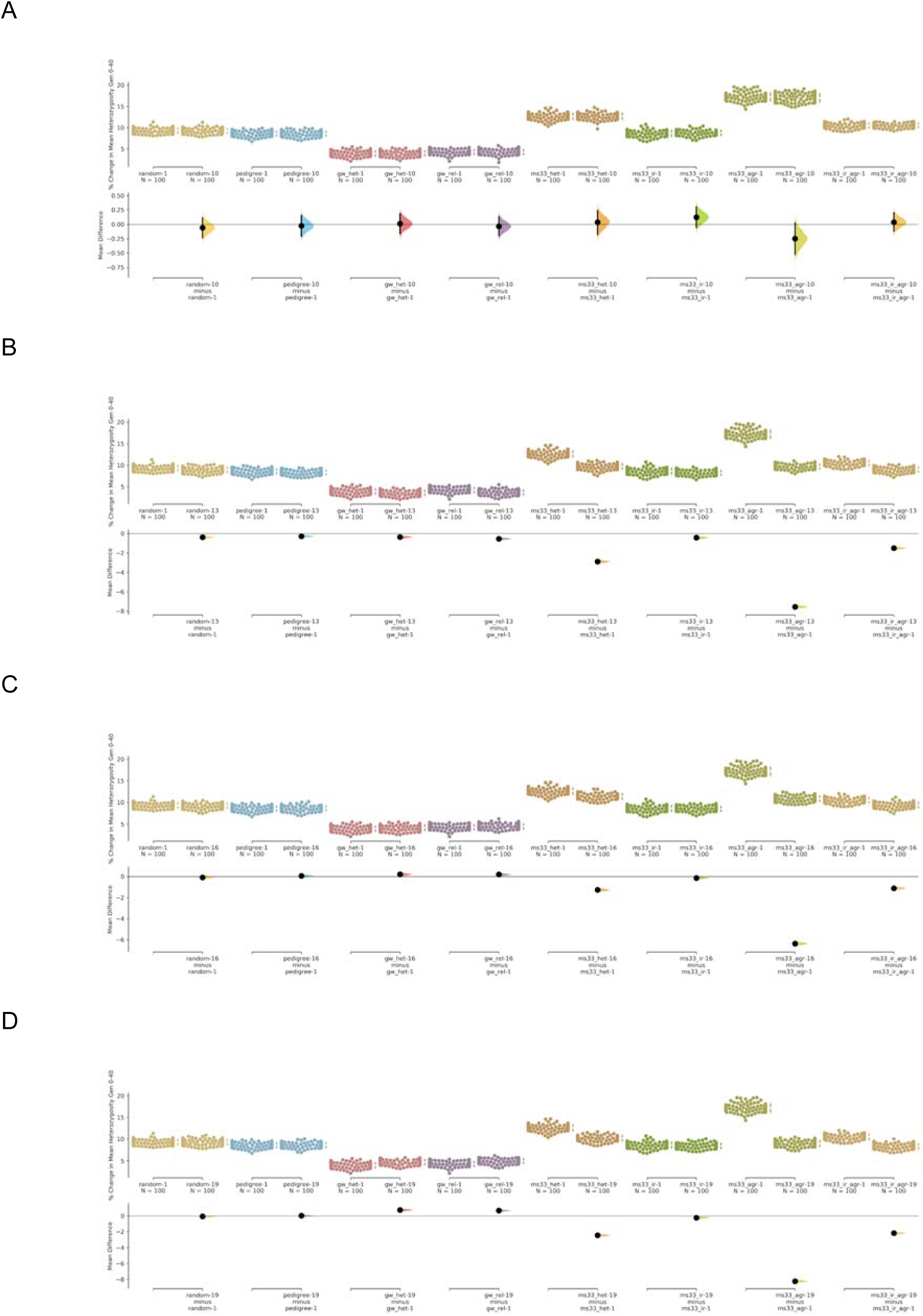
Limiting the number of matings per individual improves preservation of heterozygosity in MS models. Change in loss of heterozygosity over 40 mate choice generations. Lower values correspond to better preservation of heterozygosity. A) PS1 compared to PS10 (maximum_number_of_matings 200 to 50). B) PS1 compared to PS13 (maximum_number_of_matings 200 to 5). C) PS1 compared to PS16 (mating_pool_size 50 to 25). D. PS1 compared to PS19 (mating_pool_size 50 to 10).

Additionally, we found that halving mating pool size from 50 to 25 (0.25 to 0.125 of the total population; PS16-18) and from 50 to 10 (PS-19-21), unexpectedly led to a notable reduction in the amount of genome-wide diversity loss (heterozygosity, richness, or coefficient of inbreeding) in all MS-based mate choice schema (Figure 3C,3D, Appendix S3 — Tables S6-8). Upon further inspection, we realized this result is consistent with the mating pool size acting to moderate the number of times the same individual is chosen as a parent in a single generation and that the two parameters discussed here are not as independent as we originally envisioned when designing our simulations. This may be because the “mating pool” is randomly chosen each time a first parent is sampled, such that if the same individual is chosen twice, it will have different mating pools each time. In other words, reducing the number of individuals available as mates for a given individual reduces the number of times that individuals with low mean genetic similarity *to the rest of the population* will be over represented in the next generation. However, while limiting the number of possible mates for each individual decreased the loss of heterozygosity for MS-based mate choice models, it has no impact on the random and pedigree-based mate choice models and increases the loss of heterozygosity slightly in the two genome-wide genetic models (Figure 3C, 3D), supporting the idea above that higher variance in heterozygosity when marker sample size is low drives the unexpected relationship between preservation of heterozygosity and an individual’s mating pool size.

### Genome-wide metrics improve diversity preservation

In addition to the MS-based mate selection models, we also included several models meant to capture the viability of using genome-wide metrics in general to preserve genetic diversity. These include tracking two generation pedigrees (pedigree)- to avoid matings between very close relatives, genome-wide heterozygosity (GW-HET)- to select mates which maximize heterozygosity in the offspring, and genome-wide relatedness (GW-REL)- which prefers the most distantly related individuals as mates. We find that these three models all lead to greater preservation of genetic diversity than random mating. Perhaps more importantly, these models reduce the rate of genetic drift over time (Random model), compared to the MS models which accelerate the rate of genetic drift (Figure 1).

## DISCUSSION

In this study, we used forward population genetic simulations of a model of the canine genome to investigate the efficacy of using a small genetic marker panel (e.g. a microsatellite panel) to guide mating aimed at preserving existing genetic variation in a population, when mating decisions are made at the pairwise, rather than population, level. We ran these simulations across a range of mate choice models and demographic parameterizations using a genomic model that included both genome-wide genetic markers and a set of 33 markers distributed across the genome (herein referred to as the GW and MS33 marker sets, see methods).

Most previous work on conservation management with molecular data has focused on cases where a population can be managed by selecting the entire configuration of parents for the next generation with marker assisted selection (MAS). For example, Fernandez and colleagues (*Fernández, Toro, & Caballero, 2004) demonstrated that in a single population, management programs where parental contributions are chosen to maximize either heterozygosity or allelic richness at a set of multi-allelic markers are optimal at maintaining each of those statistics, but that heterozygosity can be better at maintaining allelic richness than vice-versa. However, no prior work, to our knowledge, has addressed whether similar strategies that optimize at the level of individual mating pairs, rather than the entire population of parents, can similarly act to preserve diversity. Consistent with this prior work, our results suggest that selecting optimal mates for individuals from an entire population using heterozygosity and other kinship metrics can act to preserve genetic diversity at markers used to calculate the test statistic (for example see Figure 2).

Importantly, however, we also found that given enough generations using small panels of markers in such a mating scheme does not preserve diversity genome-wide. In fact, mate-choice models using the MS33 marker set over time led to greater loss of genetic variation compared to random mating in the form of reduced heterozygosity and allelic richness measured using the GW marker set. Lopez-Cortegano and colleagues (*López-Cortegano et al., 2019) simulated management of subdivided populations and found that using a restricted number of markers was less effective than whole genome data but still more effective than random mating. However, the density of markers in their simulations is greater than typical microsatellite panels and they acknowledge that less dense panels would likely be less effective. Nonetheless, our results are partially consistent with this result, in that the MS33-IR model does preserve genome-wide genetic diversity better than random mating (but not allelic richness) during the course of our simulations.

Our results suggest that reducing the number of times that any given individual can contribute offspring to the next generation, either explicitly in the form of the “maximum number of matings” parameter or implicitly by reducing the “mating pool size” parameter, can act to moderate the severity of diversity loss compared to the random mating model. This finding is generally consistent with theory and prior simulation work which has demonstrated that optimal management schemes to preserve genetic diversity include limiting variance in family size, in other words, ensuring that no single individual contributes disproportionately to the next generation (Miguel Toro, Silió, Rodrigáñez, & Rodriguez, 1998).

Toro and colleagues (*Miguel Toro, Silió, Rodrigañez, Rodriguez, & Fernández, 1999) demonstrated that irrespective of variance in family size MAS should lead to better preservation of diversity than using no genetic information at all. In contrast, our results suggest that in an unsupervised pairwise parental selection scheme, limited marker panels lead to substantially more diversity loss than using no genetic information at all. A general consensus in MAS of parental populations is that pedigrees should be the primary source of kinship calculations and that small microsatellite panels are generally only useful to supplement pedigrees (M. A. Toro, Fernández, & Caballero, 2009). Our results from pairwise parental selection are consistent with this, as we have shown that using only shallow pedigrees to minimize loss of genetic diversity is preferable to using a small panel of genetic markers alone.

Genetic drift comes from two primary sources in a diploid population: variation in genetic contribution between individuals in a population and variation in genetic diversity at a given locus within an individual (J Wang & Hill, 2000). Here, we have shown (Figure 2) that the added loss of genetic variation in our simulations relative to random mating is due to accelerated loss of diversity throughout the majority of the genome that is untagged by MS33 markers. While we did not specifically explore the causes of this difference, we suspect that even in our simulations which reduce the contributions of any given individual to the next generation, that groups of individuals which happen to be most distantly related to all other individuals in a given generation across the MS33 marker set are disproportionately chosen as mates for the next generation. This variance in contribution of families (as measured by the MS33 set) across generations will act to consistently reduce effective size at markers unlinked to the MS33 marker set.

Finally, for comparison we simulated several GW mate-choice models and found, by and large, that using genome-wide genetic data to monitor genetic diversity and make mate selection decisions is far superior to small marker panels, and typically, random mating. This result has important implications for the preservation of domestic dog breeds. Most academic effort in the field of genetic diversity management over the past few decades has primarily been focused on optimal management for small populations of conservation concern where mating in the entire population can be controlled (Ballou & Lacy, 1995; Fernández, Toro, & Caballero, 2001, 2004; Kettunen, Daverdin, Helfjord, & Berg, 2017; López-Cortegano et al., 2019; Sonesson & Meuwissen, 2001). Similarly, in livestock, conservation of genomic diversity in combination with genomic selection can occur at the level of entire herds or regional populations, though also suffers from geographic partitioning and localization of conservation efforts (Bosse et al., 2015; Bruford et al., 2015; Herrero-Medrano et al., 2014; Ramljak et al., 2018; Zhao et al., 2019). In contrast, dog breeds, as well as breeds in other companion species such as cats and horses, have populations which are typically maintained by networks of individual breeders. Therefore, it is very important to understand the long term impact of different types of breeding practices in these systems.

Here, we have shown that using small panels of molecular markers is no substitute for quality pedigree information, or more importantly whole genome characterization of genetic diversity using dense genetic markers or whole genome sequence data. Our results suggest that optimal management of unsupervised companion animal populations should 1) include strictly limiting individual and family contributions to the next generation and 2) the selection of mating pairs to minimize inbreeding in offspring using deep pedigree information or, more optimally, using dense genotype data to maximize heterozygosity/minimize inbreeding in offspring, as pedigrees are often incomplete and do not incorporate variance in inheritance of IBD segments amongst related individuals (Cassell, Adamec, & Pearson, 2003; Hill & Weir, 2011; Keller, Visscher, & Goddard, 2011).

We note that we did not directly compare our results to whole-population management schemes, as such management strategies are not currently feasible for most companion animals. We suspect that such schemes will be generally superior to the unsupervised mating methods examined here, as they are better able to optimize contributions from individuals and choose the optimal (or near optimal) configuration of pairwise matings to preserve existing genetic diversity.

Most companion animal species remain relatively unmanaged with respect to genetic diversity at the breed level. As such, genetic diversity has rapidly decayed in many breeds over the past century (Jansson & Laikre, 2018). While we have not focused on optimizing the use of whole genome molecular data to preserve genetic diversity in this study, future species-specific analyses should aim to develop specific recommendations to individual breeders. For example, more realistic (non-Wright-Fisher) models would better reflect the breeding practices used in companion animal breeding. Further, in our whole genome mate choice methods we have focused primarily on maximizing heterozygosity, but preservation of allelic diversity is also an important metric to optimize, as the number of unique alleles creates a limit on the maximum heterozygosity attainable (Fernández, Toro, & Caballero, 2004). Finally, we have not considered here the needs of diversity management schemes to also consider balancing the goals of preserving genetic diversity with simultaneously eliminating deleterious variation from a population. In particular, due to the lack of past management, many companion animal breeds carry high effect deleterious mutations, and care must be taken to purge such variation without reducing linked neutral variation (Fernández et al., 2004; Hedrick & Garcia-Dorado, 2016).

Eventually, companion animal breeding may benefit from large-scale participation in databases and services aimed at tracking breed-wide whole genome genetic diversity, including awareness of adaptive and deleterious variation, to limit variance in family contributions, maximize the inclusion of genetic variation in subsequent generations, and purge deleterious variation over time. Experimentation and optimization of such a system applied to breeds in a large and diverse species such as domestic dogs would provide critical case studies to the conservation genetics community (Shafer et al., 2015), help breeders and breed organizations understand the limits of truly closed breeding, and better conserve some of the world’s most precious animal resources.

## DATA ARCHIVING STATEMENT

Data and code for this study are available at: to be completed after manuscript is accepted for publication.

## ACKNOWLEDGEMENTS

We thank the customers of Embark, who’s participation and curiosity made this work possible. We also thank our colleagues at Embark (in particular Alison Ruhe and Tiago Antao for several helpful discussions), the Embark scientific advisory board, Cornell University, and the Kevin M. McGovern Family Center for Venture Development in the Life Sciences, for their guidance and encouragement. This study was funded by Embark.

