## Appendix 2 for "Examination of the efficacy of small genetic panels in genomic conservation of companion animal populations"

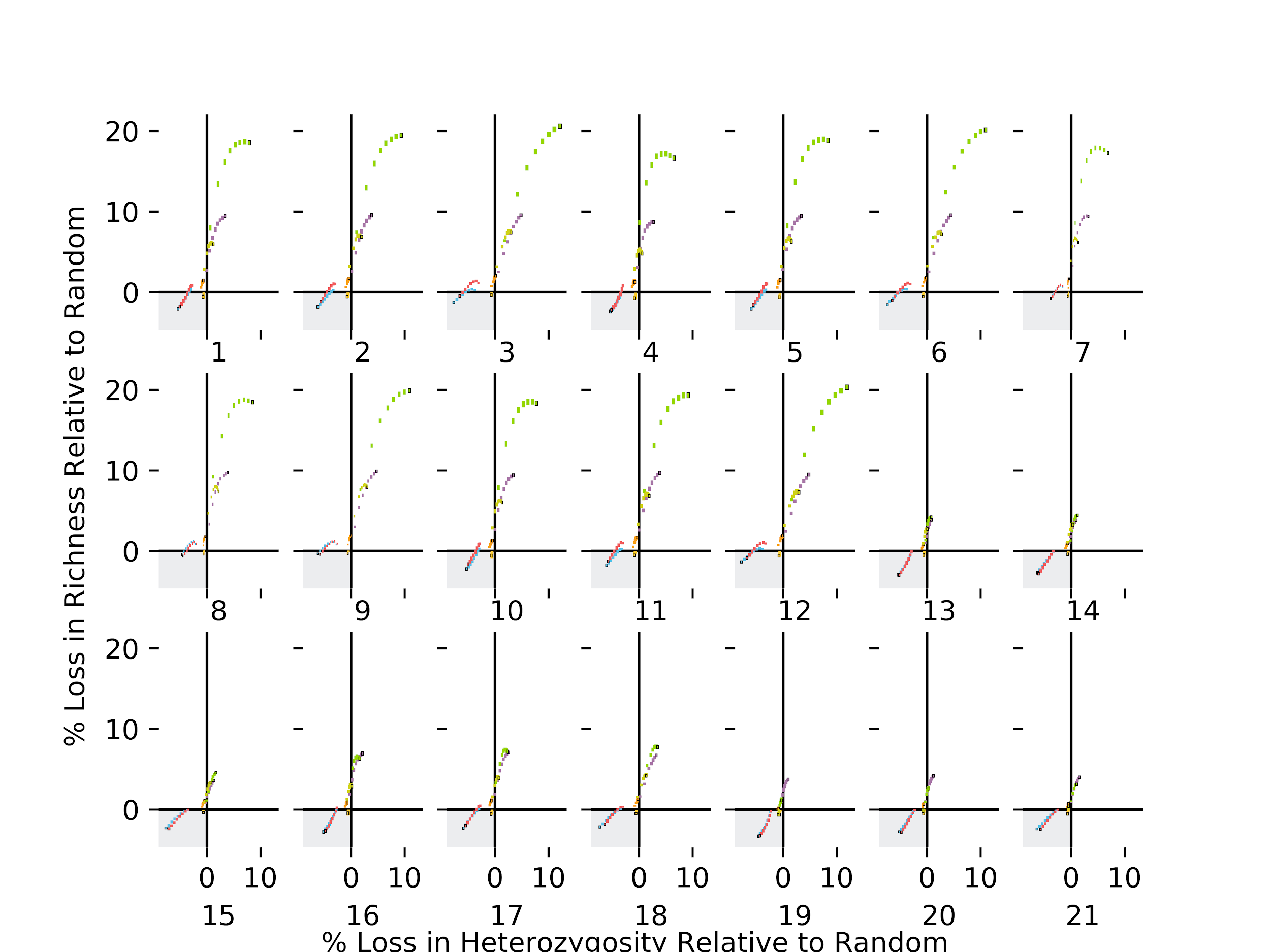

**Figure S1. Percent loss in heterozygosity and richness across all simulations**

Parameter set key is as in Figure 1.

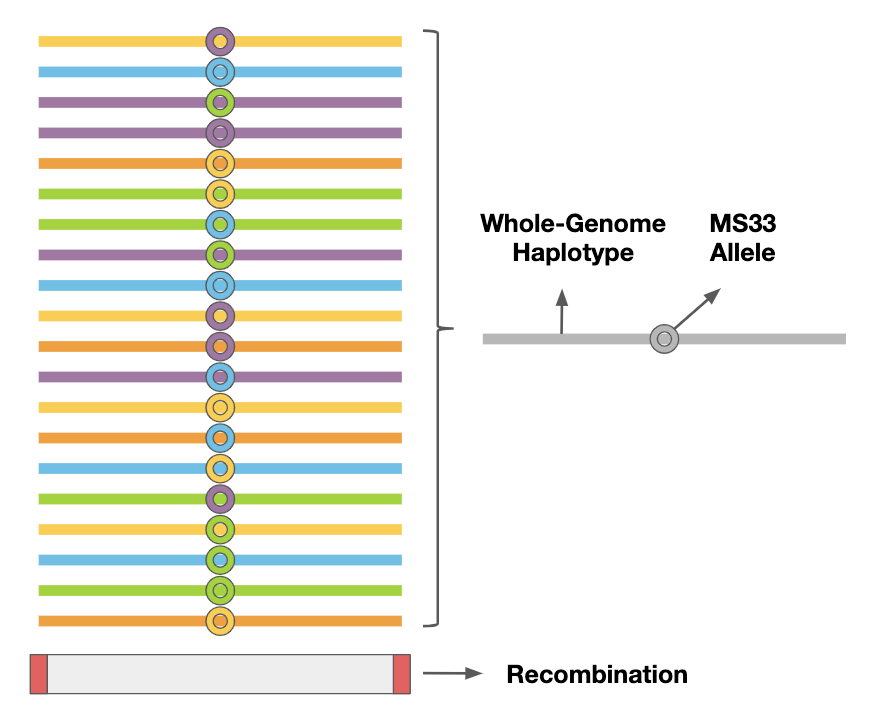

**Figure S2. Illustration of initialization state at a haplotype locus carrying a microsatellite marker**

Bars represent whole genome haplotype markers. Circles represent microsatellite markers. Colors represent alleles. Each whole genome haplotype marker is bounded by points of recombination.

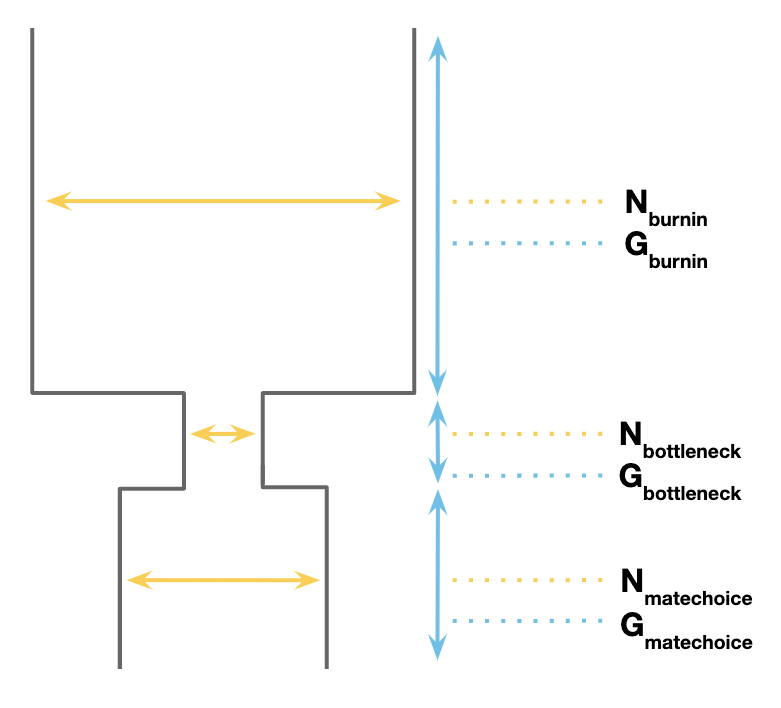

**Figure S3. Illustration of demographic model**

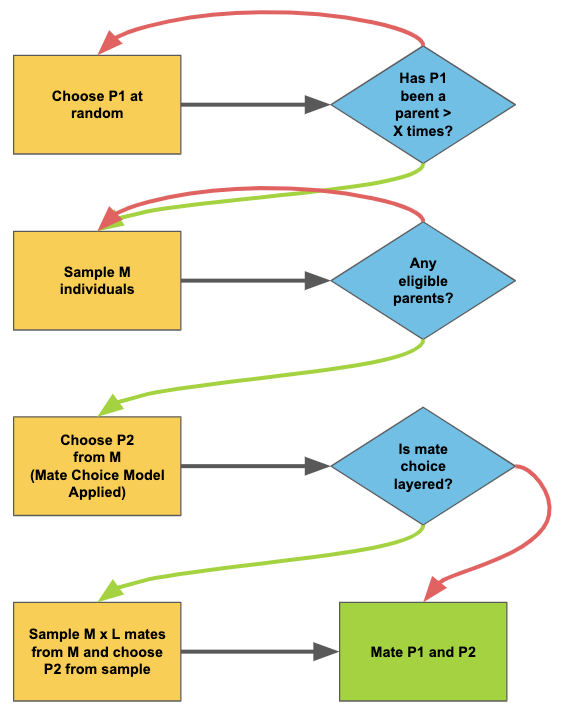

**Figure S4. Flow chart of mate choice life cycle**

Yellow boxes represent sampling events, blue diamonds represent decision points, green box represents final state (both parents chosen).

P1 — Parent 1

P2 — Parent 2

M — Random sample of individuals with size = MATING_POOL_SIZE

X — Count of number of times within generation that P2 has been used as a parent (restricted to <= MAXIMUM_NUMBER_OF_MATINGS)

L — Proportion of mating pool to sample from in layered mate choice schemes = PROPORTION_OF_MATES_FOR_LAYERED_MATE_CHOICE
